# Activity-induced fluidization of arrested coalescence in fusion of cellular aggregates

**DOI:** 10.1101/2021.02.26.433001

**Authors:** Steven Ongenae, Maxim Cuvelier, Jef Vangheel, Herman Ramon, Bart Smeets

## Abstract

At long time scales, tissue spheroids may flow or appear solid depending on their capacity to reorganize their internal structure. Understanding the relationship between intrinsic mechanical properties at the single cell level, and the tissue spheroids dynamics at the long-time scale is key for artificial tissue constructs, which are assembled from multiple tissue spheroids that over time fuse to form coherent structures. The dynamics of this fusion process are frequently analyzed in the framework of liquid theory, wherein the time scale of coalescence of two droplets is governed by its radius, viscosity and surface tension. In this work, we extend this framework to glassy or jammed cell behavior which can be observed in spheroid fusion. Using simulations of an individual-cell based model, we demonstrate how the spheroid fusion process can be steered from liquid to arrested by varying active cell motility and repulsive energy as established by cortical tension. The divergence of visco-elastic relaxation times indicates glassy relaxation near the transition towards arrested coalescence. Finally, we investigate the role of cell growth in spheroid fusion dynamics. We show that the presence of cell division introduces plasticity in the material and thereby increases coalescence during fusion.

## 1 Introduction

A general understanding of the rheological properties of multicellular tissues is important to gain insight into the physics of morphogenetic processes during development. Furthermore, robust models of these materials allow for the design and characterization of generic unit operations, such as aggregation, dispersion and fusion, which are used for the production of artificial tissues. Given its analogy to the merging of two liquid droplets, the fusion of tissue spheroids has received considerable interest as a model for soft tissue rheology. An analytical expression of the onset of coalescence of two equal viscous droplets under influence of their surface tension was first derived by Frenkel [1] and was improved and extended upon such that the dynamics of complete coalescence could be accurately modeled [2, 3]. Furthermore, various extensions to this framework have been proposed to take into account specific properties of multicellular materials, such as differences in spheroid size [4], and the presence of biological processes such as proliferation, differentiation and apoptosis, which may conflict with the assumption of conservation of mass [4, 5].

However, when using the liquid model to estimate the effective surface tension, a discrepancy emerges when comparing the fusion process with nano-indentation experiments [6]. Furthermore, in fusion assays with treatments such as Rho kinase inhibitor, it was observed that fusion may be arrested [7, 8]. This effect cannot be accounted for by a liquid material model. As such, Oriola et al. proposed to model spheroids as visco-elastic materials instead of simple liquid droplets [9].

Computational approaches such as individual cell-based models allow for the characterization of tissue-scale rheological behavior as a function of cell-scale mechanical properties [10, 11, 4, 12, 13, 14]. Using individual cell-based computational models, it was demonstrated that a divergence of the visco-elastic relaxation time can be observed within a transition region of arrested fusion, indicative of a jammed system [9]. The existence of arrested fusion could thus be traced back to the build up of elastic energy during fusion, similar to the mechanical response of tissue spheroids in parallel plate compression experiments [15].

In this work, we derive a simplified analytical expression for the coalescence of visco-elastic tissue spheroids, obtaining similar results to [9]. We demonstrate the applicability of this expression by using an individual-cell based model approach to simulate the fusion process and, using this expression, we are able to compare the relaxation dynamics of fusion to the characteristic timescales involved in the preceding aggregation (or compaction) process in which the individual tissue spheroids have been formed. Finally, we extend this simulation framework to include a morphological model of the cell cycle. In analogy to active motility, cell division and growth may introduce excitation that may induce fluidization, as was observed in other systems [16, 17, 18]. Here, we investigate whether in the framework of spheroid fusion, there is an additional active contribution of cell division beyond a mere correction for the increase in volume, and assess to what extent this contribution may unjam the cellular material [4, 5] and recover tissue coalescence.

## 2 Material and Methods

### 2.1 Individual cell-based model

We follow an individual cell-based model approach as described in detail in Smeets et al. [19, 20]. This model has already been used to study cell aggregation and compaction dynamics and is here extended to take into account the cell cycle to simulate during the fusion of two tissue spheroids. In brief, the model is based on a simulation framework introduced by Delile et al. [21], where cells are simulated as self-propelled particles. Conservative active forces are exchanged between neighboring cells, similar to [9]. The connectivity network is based on a Delaunay triangulation of the cell center coordinates. For the edges of this network, a symmetric central potential is calculated, which is parameterized by adhesion *w*_*a*_ and cortical tension *w*_r_. Protrusive active forces are responsible for active migration with velocity *v*_*t*_. These forces are calculated in the direction of the cell’s polarization which is randomly diffusing with rotational diffusivity *D*_*r*_. Hence, activity from cell motility may be parameterized as 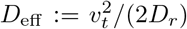. Overdamped equations of motion are integrated to evolve the system over time. Finally, cells are able to growth and multiply based on a cell cycle model, which is explained in (SI 1). The model parameters are listed in tables in (SI 2).

### 2.2 Simulation setup

The simulation pipeline is shown in Fig. 1 and follows a sequence of generic unit operations to form a small tissue via fusion. In the first step we simulate the seeding and spontaneous aggregation of two aggregates for 24 hours, each in their own micro-well, similar to [20]. This guarantees that the shape of each spheroid is consistent with respect to its underlying mechanical properties. It should be noted that, depending on the mechanical properties that govern the aggregation process, this relaxed configuration may have a highly irregular shape. Any (few) remaining cells that did not get incorporated in the main aggregates, are removed from the simulation in analogy to a real fusion experiment. In the second step, the two aggregates are transferred to a larger micro-well and are brought into contact with each other, simulating another 50 hours during which the fusion process naturally progresses. In the final step, we extract the spheroid contours and fit two circles to compute the contact angle *θ* for each simulation, as explained in (SI 3). Each realization of a simulated fused spheroid is initialized independently. To calculate the average fusion dynamics, i.e. the average of sin(*θ*(*t*))^2^ across all repeats, we take into account that not all spheroids start fusing at the same time, because the initial contact between the spheroids can be weak. To account for this, we define the time at which fusion starts as the last time at which we observe two distinct objects in the extracted contours. For further analysis, the extracted shape measures are shifted in time using this offset.

**Figure 1:**
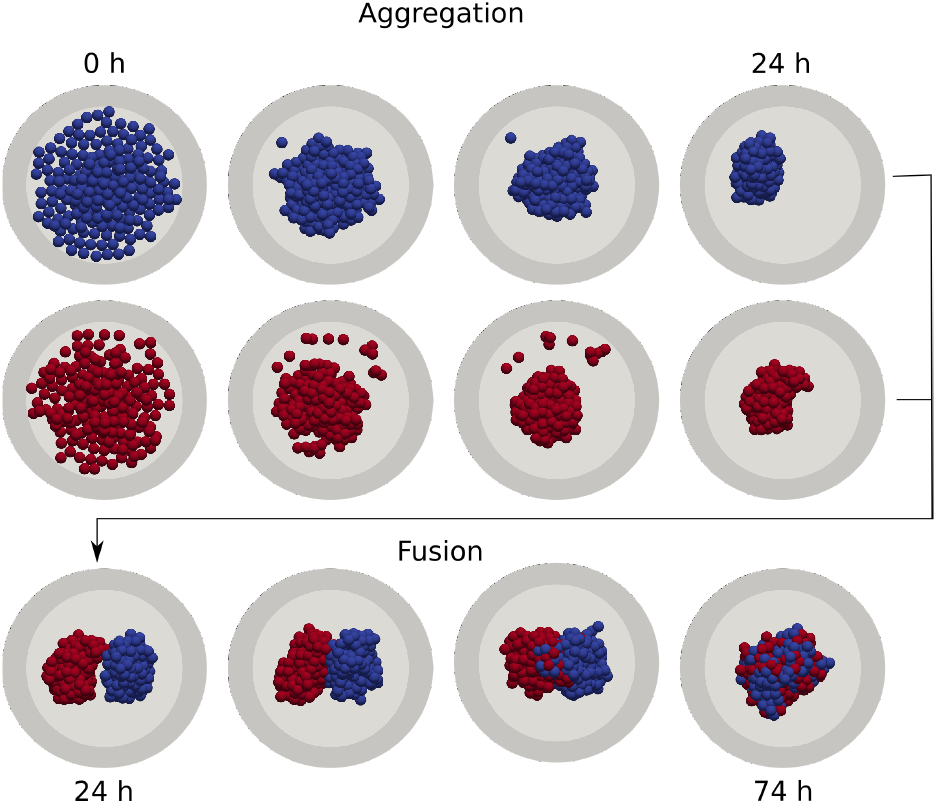
Individual cell-based model of cell aggregation and spheroid fusion: Gravitationally deposited cells aggregate in independent cylindrical wells for 24h top). Next, two aggregates are extracted and combined in a single fusion simulation, which continues for a further 50h (bottom).

### 2.3 Visco-elastic approximation of the fusion process

The fusion dynamics of two equal visco-elastic spheres can be approximated by the differential equation

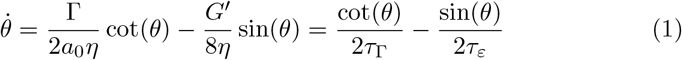

as derived in (SI 4). Here, *θ* is the angle as shown in Fig. 2, and is influenced by the surface tension Γ, the radius of the spheroid before fusion *a*_0_, the apparent viscosity of the tissue *η* and the shear modulus of the tissue *G*^’^ We combine these parameters into two characteristic time constants; *τ*_Γ_ is the visco-capillary time, and *τ*_*ε*_ is the visco-elastic time. The analytical solution of Eq. (1) is

**Figure 2:**
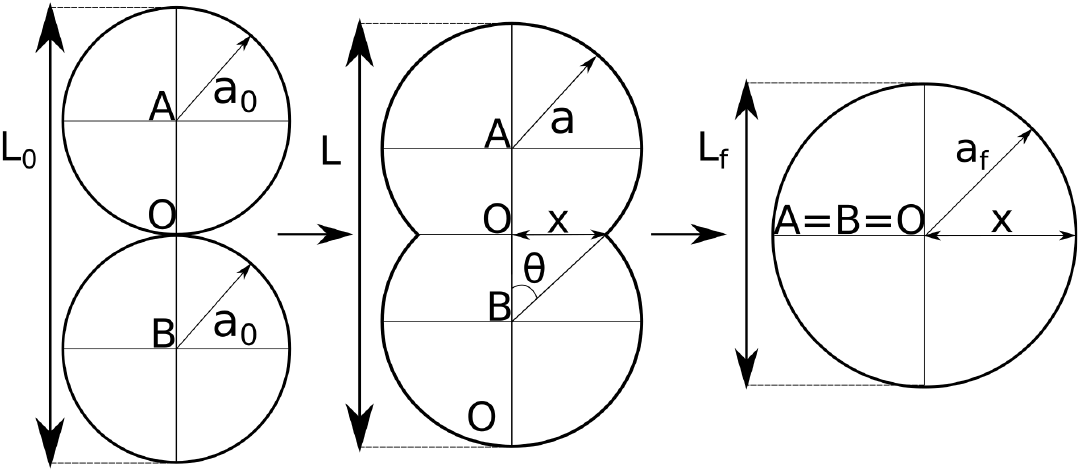
Shape evolution during spheroid fusion adapted from [3]

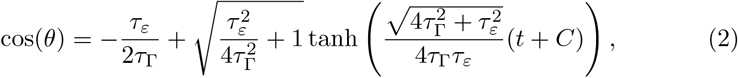

in which the complex number *C* can be obtained from the initial conditions *θ*(*t* = 0) = *θ*_0_

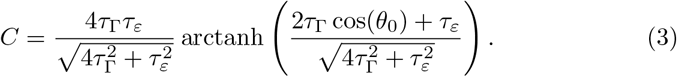

In analogy to previous studies on spheroid fusion, the fusion dynamics are reported as sin^2^(*θ*) = (*x/a*)^2^ with *a* the radius of the spheroid and *x* the contact radius which both vary in time [11, 4, 12, 13, 9]. Other studies perform the analysis based on (*x/a*_0_)^2^ [5, 6, 14, 10], but this is less consistent with the underlying theory (see SI 5). Eq. (2) is used to calculate sin^2^(*θ*). When fitting, we use this equation with free variables: *τ*_Γ_, *τ*_*ε*_ and *θ*_0_. In order to compare to simulated fusion experiments, *θ*_0_ is retained as a free variable since the discrete initial adhesion between the first contacting cell pair will permit a rapid relaxation towards a non-zero initial angle, *θ*_0_. Based on the fitted values, the predicted equilibrium angle *θ*_*eq*_ can be obtained as

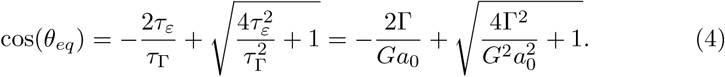

## 3 Results

### 3.1 Arrested coalescence dynamics

The average dynamics of simulated tissue spheroid fusion are consistent with the derived visco-elastic material model, expressed in the time evolution of contact angle *θ*, Eq. 1, as shown in Fig. 3B where we varied the cell activity *D*_eff_. At sufficiently low *D*_eff_, coalescence appears arrested and complete fusion is not attained. Given the correspondence to the visco-elastic model, the parameters *τ*_Γ_ and *τ*_*ε*_ can be interpreted as characteristic timescales of the multi-cellular visco-elastic material. From the fit of *τ*_Γ_, *τ*_*ε*_ and the instantaneous initial angle *θ*_0_, we are able to calculate the equilibrium fusion angle, *θ*_*eq*_, using Eq. (4). When varying the cell activity *D*_eff_ and the repulsive energy *w*_r_, two distinct regions can be recognized based on *θ*_*eq*_, Fig. 3F. For low activity (*D*_eff_) fusion is arrested, while at higher activities fusion is complete. Higher levels of repulsion *w*_r_ require more cell activity to fluidize the material and hence to attain complete fusion. Similarly, the characteristic visco-capillary time *τ*_Γ_ increases when cell activity decreases and when cell repulsion increases, Fig. 3E. When reducing cell activity within the fluidized region, the visco-elastic time *τ*_*ε*_ gradually increases and displays a divergence near the transition line towards arrested coalescence. This divergence of elastic relaxation time is indicative of an underlying glass transition, as was already pointed out in [9], although based on somewhat different analytical and computational models. The transition from arrested to complete fusion as characterized by *θ*_*eq*_, coincides with the glass transition as characterized by the divergence of *τ*_*ε*_, as shown in Fig. 3D. Since this computational model is based on the same individual cell-based framework that was used to simulate the aggregate formation process, we are able to make a direct comparison between dynamics of aggregation and dynamics of tissue spheroid fusion. At sufficient cell activity, the aggregation process is consistent with the dewetting of a liquid film from a surface, as was demonstrated in [20]. Upon decrease in activity, this correspondence is lost. Instead, the aggregate density *ρ*_*a*_ follows the dynamics of granular compaction, characterized by stretched exponential relaxation (KWW law):

**Figure 3:**
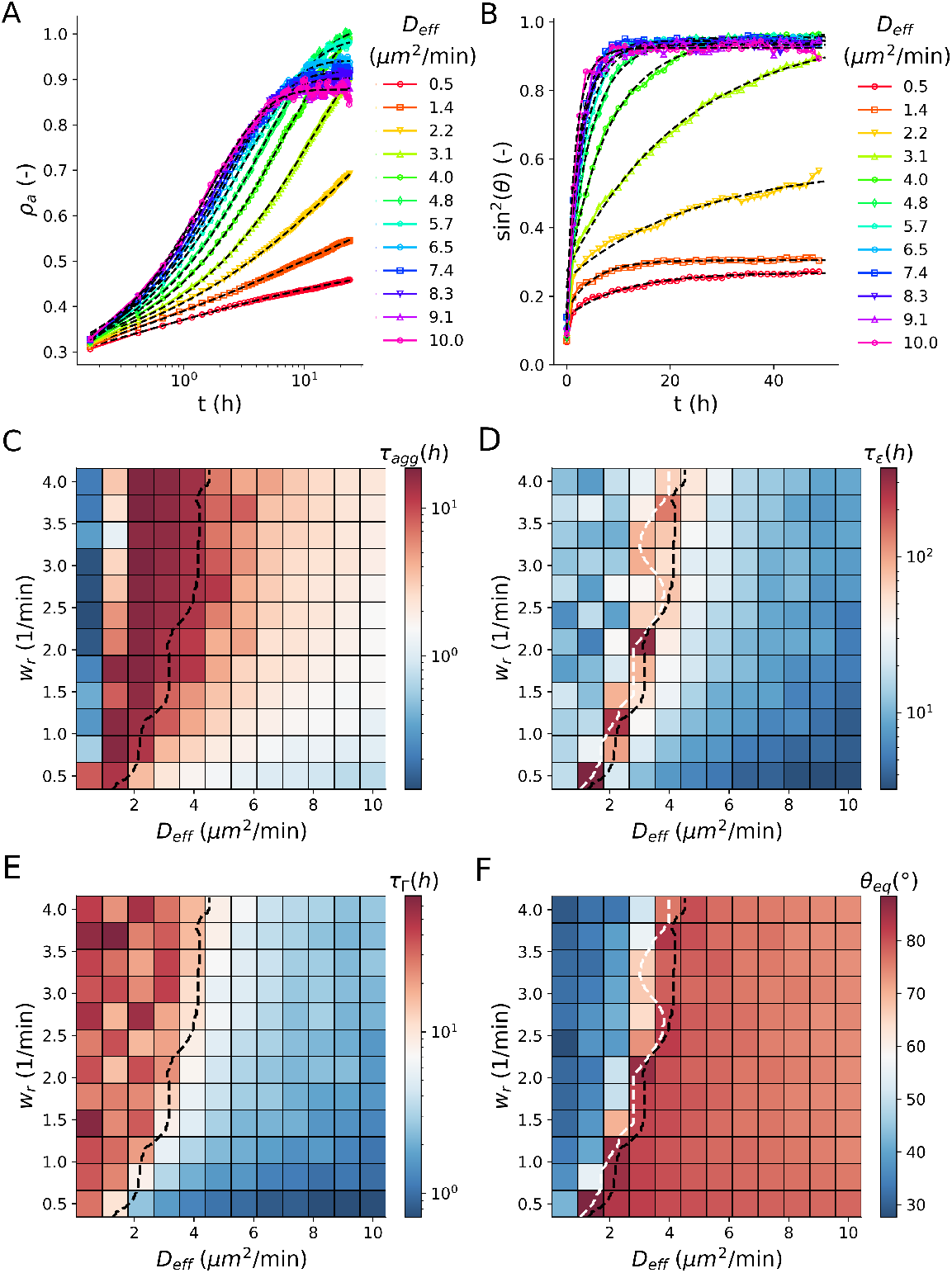
**(A)** Average (N=100) apparent density *ρ*_*a*_ in function of time for varying activity *D*_eff_ at constant *w*_r_ = 2.09 min^−1^. The KWW law fits, Eq. (5), are shown as black dashed lines. **(B)** Average (N=50) fusion dynamics, represented as sin^2^(*θ*), for varying cell activity *D*_eff_ at constant *w*_r_ = 2.09 min^−1^. The simulated data is fitted by calculating sin^2^(*θ*) based on the solution for *θ* Eq. (2, 3). **(C)** Estimated values of *τ*_*agg*_ based on KWW law Eq. (5) during aggregation for varying cell repulsion *w*_r_ and cell activity *D*_eff_. **(D)** to **(F)** Estimated values of the visco-elastic time constant *τ*_*ε*_, the visco-capillary time constant *τ*_Γ_ based on fitting fusion dynamics in sin^2^(*θ*) using Eq. (2, 3), and the equilibrium angle *θ*_*eq*_ calculated using Eq. (4). The black dashed line shows the separation of arrested versus complete fusion based on *θ*_*eq*_. The white dashed line represents the separation between two regions of low *τ*_*ε*_. Both lines are obtained by performing a watershed segmentation on the images of *θ*_*eq*_ and log(*θ*_*ε*_), respectively.

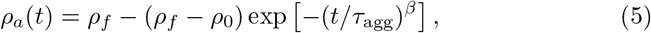

with initial density *ρ*_0_, final density *ρ*_*f*_, exponent *β* and characteristic timescale *τ*_agg_, see Fig. 3A. During compaction, a similar glass transition as observed during the fusion process can be recognized from the divergence of *τ*_agg_ [20]. For the same underlying cell properties *D*_eff_ and *w*_r_, these two transitions closely align, as demonstrated in Fig. 3B, although the transition in the fusion process consistently occurs at higher values of cell activity. We hypothesize that this discrepancy is due to the additional constraints in cell movement imposed by the geometry of the spheroid during the fusion process, compared to the loose network connectivity that characterizes the aggregation phase. Due to the sequential simulation of aggregation and fusion, which mimics the experimental procedure, additional compaction may occur during fusion, particularly at low activity when compaction proceeds slowly. We verified that the effect of this additional compaction on the predicted equilibrium angle *θ*_*eq*_ is minor, as shown in SI Fig S7, where we compare fusion of self-assembled aggregates to artificially compacted spheroid structures.

### 3.2 Increase of coalescence due to cell division

Next, we turn to the role of cell division in the arrest of coalescence. For this, we simulated the fusion dynamics in the presence of cell proliferation, see SI 1. Although cell proliferation is not explicitly accounted for in the derivation of our analytical model, Eq. (1) still fits well the fusion dynamics of our simulated spheroid fusion in the presence of cell division, Fig. 4A. Fig. 4B and C compares the characteristic visco-elastic time constant *τ*_*ε*_ and the predicted equilibrium angle *θ*_*eq*_ without (*k*_div_ = 2 *×* 10^−8^ h^−1^) and with (*k*_div_ = 0.06 h^−1^) cell division, when varying cell motility *D*_eff_ at repulsive energy *w*_r_ = 2.09 min^−1^. This enables us to evaluate the effect of cell growth / division as an additional source of biological excitation compared to active cell motility. The simulated cell division rate corresponds to a cell cycle period of approximately 16.7 hours, which is lower than many commonly used cell lines which are used for spheroid fusion. Yet, even at this relatively high division rate, we do not see a strong shift in critical activity beyond which the material appears fluid-like, as indicated by the coincidence of the peak in *τ*_*ε*_ for varying *D*_eff_ with and without cell division (Fig. 4B). However, we do observe that the presence of cell division greatly increases the overall visco-elastic relaxation time, indicating a decrease in the apparent elasticity of the multi-cellular material. Furthermore, cell division markedly increases the equilibrium angle *θ*_*eq*_ in the arrested fusion region. Hence, in the simulated configuration, cell division appears to recover coalescence by increasing the plasticity of the tissue. However, it has no strong influence on the location of the fluidization transition.

**Figure 4:**
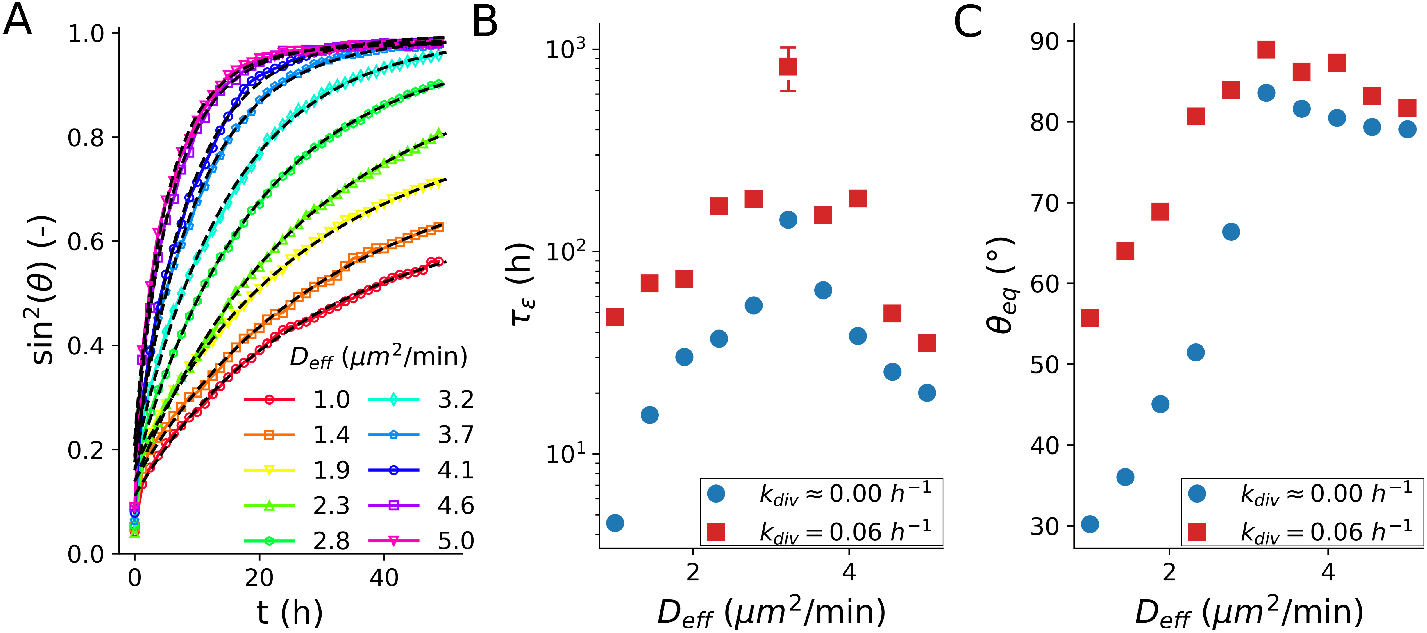
**(A)** Average (N=100) fusion dynamics with simulated cell cycle, quantified as sin^2^(*θ*), during 50 hours for varying cell activity *D*_eff_ at constant *w*_r_ = 2.09 min^−1^ and *k*_div_ = 0.06 h^−1^. The simulated data is fitted by calculating sin^2^(*θ*) based on the solution for *θ* Eq. (2, 3). **(B)** to **(C)** Comparison of fusion dynamics in presence and in absence of cell division, as a function of cell activity *D*_eff_ at a constant *w*_r_ = 2.09 min^−1^. The estimated values for the visco-elastic time constant *τ*_*ε*_ are based fitting fusion dynamics in sin^2^(*θ*) using Eq. (2, 3), the predicted equilibrium angle *θ*_*eq*_ calculated by Eq. (4).

## 4 Discussion

In this work, we derived an expression for the arrested coalescence of tissue spheroid fusion, based on visco-elastic material properties. Simulations of a minimal individual cell-based model of the fusion process showed that this expression is able to describe the transition of liquid-like to arrested coalescence dynamics. Furthermore, a divergence in the visco-elastic relaxation time indicates the presence of jammed or glassy relaxation behavior near the transition towards arrested coalescence. These findings are highly similar to recent work from Oriola et al. [9], hence a brief comparison between these contemporary results is appropriate. First, the analytical expressions for the dynamics of *θ* (Eq. (1) in [9], compared to Eq. 1) are based on somewhat different assumptions. Our model is an extension of the model of Pokluda [3] by adding an elastic energy term in the equation which was suggested by [22, 23]. This has the advantage that in the absence of elasticity, the model of Pokluda is retrieved. The downside of our approach is that this leads to an inconsistency in the strain and strain rate for viscous energy dissipation rate and the elastic energy rate (we note this difference as *ϵ* and *ε*). For the viscous dissipation term we have according to Pokluda [3] 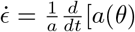. On the other hand, the strain in arrested coalescence is suggested to be *ε* = 1 *− a*(*θ*)(1 + cos(*θ*)*/*(2*a*_0_) [22, 23]. Therefore the strain rate is 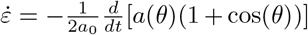. These expressions are equivalent in magnitude only for the onset of fusion i.e. *θ ≈*0 and *a*(*θ*) = *a*_0_. In contrast to our hybrid model, Oriola et al. [9] consistently use the strain rate as suggested for arrested coalesence [22, 23]. Furthermore, they introduce elasticity by considering the spheroid as an incompressible Kelvin-Voigt material, instead of adding a separate elastic energy term, hence obtaining a critical value for the Young’s modulus for which fusion is inhibited. Moreover, whereas they include a ‘pre-strain’ to account for the initial fusion onset, we allow for an instantaneous initial angle *θ*_0_. Still, both models are parameterized by two essential characteristic timescales: the visco-capillary timescale *τ*_Γ_ and the visco-elastic timescale *τ*_*ε*_. Secondly, one key difference between the individual cell-based model in this work and the one in [9] is the implementation of the active cell motility. Their model is based on ‘protrusions’ defined on the level of cell-cell bonds, and effectuates persistence by means of a protrusion lifetime per bond. Our model is based on a polarization direction defined for each cell which diffuses over time. Nonetheless, the similarity of the main results underpins that the transition of liquid-like to arrested coalescence is a generic phenomenon which is not dependent on the precise assumptions of the visco-elastic description, nor on the implementation details of the individual cell-based model.

The glassy relaxation dynamics during fusion mirror the dynamics of the aggregation process during which the initial tissue spheroids are formed. A direct comparison between these two unit processes shows that there is a clear correspondence between, on the one hand, the transition from granular compaction to liquid dewetting during the aggregation phase, and on the other hand, the transition from arrested coalescence to liquid behavior during the fusion phase. However, this transition occurs for slightly smaller values of cell activity in the case of aggregate formation. Experimental confirmation of this correspondence can be found in studies involving Rho kinase inhibitor, which has been observed to cause arrested dynamics during aggregate formation of human periosteum-derived cells [20], as well as inhibit fusion of spheroids from human mesenchymal stem cells [8] and of embryonal chicken organoids [7].

In addition, we considered the effect of cell division on the dynamics of the fusion process. In the framework of arrested coalescence, we showed that the presence of cell division may recover coalescence of fusion. Other studies on the role of cell growth in biological active matter systems, for example in simulations of two-dimensional epithelial tissues [18], or in growing 3D cell aggregates [16], observed an increase of fluidization induced by cell division. More generally, an overview of the role of cell division in tissue rheology and mechanics is provided in [24]. However, in our simulations, the effect of cell division on the location of the fluidization transition was limited, at least for realistic cell division rates compared to the timescale of fusion. Instead, cell division appeared to increase plasticity of the glassy material and thereby improve coalescence in the arrested phase. However, it should be noted that the absence of fluidization as a result of cell division in simulations could be partly due to the minimal representation of cell shape, which introduces artificial energy barriers and thereby overly penalizes neighbor exchanges. This shortcoming could be addressed in more detailed tissue models, such as vertex models [25] or deformable cell models [26, 27].

In practice, technologies that involve the production of artificial tissues frequently incorporate subsequent steps of micro-aggregation and tissue assembly, where the latter often relies on the (partial) fusion of spheroids to create larger tissue constructs [28]. To complicate matters, all these steps are typically accompanied by biological processes such as cell division, production of extracellular matrix, cell differentiation or apoptosis. However, since the physical description of the underlying aggregation and fusion dynamics is highly generic, each of these steps may be parameterized in terms of its characteristic material properties, allowing for the comparison within and between different culture conditions and production formats. As such, continued efforts towards the characterization of structure, rheology and mechanics of these artificial tissues will become indispensable.

## Author Contributions

B.S. and H.R. conceived the project, S.O. and B.S. designed and conducted the simulations, S.O. performed the mathematical analysis, S.O. performed data analysis. S.O., M.C., J.V. and B.S. wrote the manuscript. All authors commented on the manuscript.

## Acknowledgments

We thank Jiř íPešek for proofreading the derivations.

