## Supplementary Materials for "Activity-induced fluidization of arrested coalescence in fusion of cellular aggregates"

February 26, 2021

#### 1 Cell cycle

In this section we discuss how the stationary distribution of cell radii from a stochastic growth model can approximately reproduce the experimental distribution of cell sizes, as obtained in [1]. Thus, when starting the simulation based on samples of this stationary distribution, the cell population will not drift. This ensures that we only observe the effect of the growth and division, and their excitation of the system, and not a change of the cell population. More information on the implementation of the growth phase and the division phase in our simulation framework can be found in [2].

Growth of the cell is modeled using the Qu growth model [3] which states that the radius of a cell increases during the growth phase according to

$$\dot{r} = K_c \left(\frac{r_0}{r}\right)^2. \tag{1}$$

$K_c$ is a specific growth constant [2]. After integration and using the initial condition that $r(t=0) = r_{\text{init}}$ we find that

$$r = \left[3K_c r_0^2 t + r_{\text{init}}^3\right]^{1/3}. \tag{2}$$

This is a more general case in which the cell has already grown from $r_0$ to $r_{\text{init}}$. Over a time $T_g$ the cell grows from the radius $r_0$ to its maximal radius $r_{\max}$, after which cell division should start. Based on this the growth rate constant $K_c$ is calculated.

$$K_c = \frac{1}{3T_g r_0^2}\left(r_{\max}^3 - r_0^3\right). \tag{3}$$

Division splits the cell in two equal volumes, therefore $r_0 = 2^{-1/3} r_{\max}$, resulting in

$$K_c = \frac{r_{\max}}{2^{1/3}\, 3T_g r} \tag{4}$$

This expression determines the growth rate of each cell in our simulation and Eq. (1) is numerically integrated. A heterogeneous population is simulated by using that $r_{\max} \sim \mathcal{N}(\bar{r}_{\max}, \sigma^2_{r_{\max}})$. for each growth phase. This means that some cells will grow to larger sizes, and, because $K_c$ is fixed, take longer to do so. Furthermore some cells will grow faster (Eq. (1)) because $r_0 = 2^{-1/3} r_{\max}$, and $r_{\max}$ is different for each cell cycle. Now, because we want this simulated population of growing cells to reflect the experimental distribution of cell radii in terms of its mean $\bar{r}$ and standard deviation $\sigma_r$, we have to determine $\bar{r}_{\max}$ and $\sigma^2_{r_{\max}}$. First, we calculate the average radius of a growing cell in terms of $r_{\max}$. Using Eq. (2) and Eq. (3) we find the closed expression for the growth cycle from $\bar{r}_0$ to $\bar{r}_{\max}$

$$r = \left[ \frac{\bar{r}^3_{\max} - \bar{r}^3_0}{T_g} t + \bar{r}^3_0 \right]^{1/3} \tag{5}$$

which allows us to calculate the average radius of a growing cell in terms of $\bar{r}_{\max}$

$$\bar{r} = \frac{3}{4} \frac{a^3 + a^2 + a + 1}{a^2 + a + 1} \bar{r}_{\max} \tag{6}$$

with $a = 2^{-1/3}$. This expression is only approximately true, so later on we will correct for it. In this model the effective growth time is also influenced by $\sigma_{r_{\max}}$, which has a small effect on the calculation of the average. The value for $\sigma_{r_{\max}}$ has to be determined numerically. We do this by simulating growth cycles with $T_g = 1$, sample rate 0.01 and simulate for $5000\, T_g$ to calculate $\sigma_r$ for this ensemble. Because an estimation of a standard deviation is inherently noisy we take the average over 1000 runs. This process we repeat for various values of $\sigma_{r_{\max}}$. By doing so we identified a zone with a minimum (Fig. 1A), we then did a second more detailed minimum search for this zone, similarly but using 2500 runs. We identified a $\sigma_{r_{\max}} = 1.481 \mu m$. Based on the simulation results at the identified $\sigma_{r_{\max}} = 1.481 \mu m$ we propose a correction factor of 0.95 on Eq. (6). The resulting distribution matches well with the experimental distribution, as demonstrated in Fig. 1B.

Having shown that this cell cycle model replicates the distribution of radii in growing cells, we initialize cells in the fusion simulations. We assume that at the start of the fusion, all cells are growing cells. It is non-trivial to generate values for the cell radius $r$ and their corresponding boundaries $r_0$ and $r_{\max}$, since these three are not independent random variables. Hence, we simulate subsequent fictive growth cycles to generate this set based on the optimized values for $r_{\max}$ and $\sigma_{r_{\max}}$.

#### 2 Model parameters

Table 1: Simulation parameters that are unique to or specifically set for the simulation of the initial settling phase. Parameters that are not indicated in this table are identical to the ones further given in Table 2.

| **Parameter** | **Symbol** | **Value** | **Units** |
|---|---|---|---|
| Tangential friction | $\gamma_t$ | 0.02 | kPa·s/m |
| Adhesive relaxation rate | $w_a$ | 0 | $\text{min}^{-1}$ |
| Relative active velocity | $v_t$ | 0 | m/min |
| Viscosity medium | $\eta_{\text{eff}}$ | 0.8 | mPa·s |
| Gravitational acceleration | $g$ | 9.81 | $\text{m/s}^2$ |
| Total medium area | $A_{\text{m}}$ | 1.8 | $\text{cm}^2$ |
| Total medium volume | $V_{\text{m}}$ | 1.0 | ml |
| Aggregation time | $t_a$ | 24 | h |
| Timestep | $\Delta t$ | 8 | ms |

Table 2: Simulation parameters used for the individual cell-based model, with indication of the source or rationale for each used value. 'Free' parameters that are studied in this work are not reported in this table.

| Parameter | Symbol | Value | Units | Source |
|---|---|---|---|---|
| Microwell radius | $R_{\mathrm{w}}$ | 108.56 | $\mu$m | Measured |
| Microwell height | $h_{\mathrm{w}}$ | 2000 | $\mu$m | Micro-well design |
| Average cell radius | $\langle r \rangle$ | 8.99 | $\mu$m | Measured |
| Standard deviation cell radius | $\sigma_{\mathrm{r}}$ | 1.42 | $\mu$m | Measured |
| Average maximum cell radius before division | $r_{\mathrm{max}}$ | 9.438 | $\mu$m | Optimized |
| Standard deviation maximum cell radius before division | $\sigma_{\mathrm{r_{max}}}$ | 1.481 | $\mu$m | Optimized |
| Number of cells | $N$ | 2·200 | - | Set |
| Cell density | $\rho_c$ | 1100 | kg/m$^3$ | [4] |
| Medium density | $\rho_m$ | 1000 | kg/m$^3$ | |
| Effective medium viscosity | $\eta_{\mathrm{eff}}$ | 100 | mPa·s | |
| Gravitational acceleration | $g$ | 9.81 | m/s$^2$ | |
| Effective contact stiffness | $\hat{E}$ | 888 | Pa | Estimated |
| Tangential friction | $\gamma_t$ | 1.0 | kPa·s/m | Reference value |
| Relative Normal friction | $\overline{\gamma}_n$ | 2.0 | $\gamma_t$ | Estimated |
| Cell-cell adhesion | $w_a$ | 4.0 | min$^{-1}$ | Estimated |
| Rotational diffusivity | $D_r$ | 0.5 | min$^{-1}$ | Estimated |
| Attractive reach | $c_{\mathrm{max}}$ | $2^{1/3}$ | - | Estimated, [5] |
| Contact area factor | $a$ | 1.3697 | - | [5] |
| Protrusion tuning angle | $\eta_{\mathrm{min}}$ | 0.15 | - | [5] |
| Relative resting distance | $c_{\mathrm{eq}}$ | $(\pi/(2\sqrt{3}))^{1/2}$ | - | [5] |
| Aggregation time | $t_a$ | 24 | h | Set |
| Fusion time | $t_f$ | 50 | h | Set |
| Timestep | $\Delta t$ | 0.25 | s | Optimized |

### 3 Shape extraction

The main geometric measures of interest for modeling the dynamics of the fusion process are the individual sphere radius $a(t)$ and the contact radius $x(t)$. To compute these, we first project the simulation on the $X/Y$-plane and extract the aggregate contour. Next, we fit two circles with equal radius on this contour. The result of this procedure is shown in Fig. 2. Since this analysis routine only needs the segmented contour of the aggregates, it is equally applicable to experimental data obtained using time-lapse microscopy. To compute the contour of the combined tissue spheroid, we start with a cell (circle) on the outside of the aggregate, and then, based on the connectivity network, progressively add the circle arcs of neighboring cells until the starting cell is encountered once more.

Next, we discretize the contour in 1000 equally spaced points along the contour. In the circle fitting procedure we search for the center points of circle A and B i.e. $\{(x_a, y_a), (x_b, y_b)\}$ for which the sum of squares of the difference between the minimal center-contour distance and the mean center-contour distance is minimized. For this we start with two guess points $\{(\hat{x}_a, \hat{y}_a), (\hat{x}_b, \hat{y}_b)\}$. We calculate the distance between these points and all contour points $i$ so that $d_{a,i} = \sqrt{(x_i - (\hat{x}_a)^2 + (y_i - (\hat{y}_a)^2}$ and $d_{b,i} = \sqrt{(x_i - (\hat{x}_b)^2 + (y_i - (\hat{y}_b)^2}$. Each contour points $i$ is assigned to the circle corresponding to its closest center point, therefore $d_i = \min\{d_{a,i}, d_{b,i}\}$. Next, we minimize the sum of squared errors $\sum_{i=1}^{n}(d_i - \bar{d}_i)^2$ with our center points of circle A and B as degrees of freedom. The radii of circle A and B are then given by $\bar{d}_a$ and $\bar{d}_b$. It has to be noted that we always fit the best two equals circles on the contour, and due to the irregular shape of aggregate the centers of these circles never coincide, thus even in the liquid limit our observable $\sin(\theta) = (x/a)^2 < 1$. This implies that the spheroid will always have a small apparent elasticity.

### 4 Derivation of visco-elastic fusion dynamics

The classical model for the coalescence of two spherical droplets [6] is not able to explain the phenomenon of arrested coalescence. It has been shown experimentally that an internal elasticity caused by structures inside the droplet is a way to arrest the coalescence in a controlled fashion [7, 8]. However, the scope of these studies was limited to explaining the equilibrium. In what follows we add an additional elastic energy term, suggested by [7, 8] to expand the current liquid framework. We start by introducing the liquid model, which has already been described in detail in literature [6, 9, 10].

In the liquid model, coalescence dynamics are the result of the balance between the work rate of surface tension and viscous energy dissipation together with the assumption of conservation of mass, which at constant density entails conservation or volume. For biological systems the latter assumption is precarious because the cells inside the spheroid might proliferate, shrink, or undergo apoptosis. However, If the coalescence dynamics are the dominating time scale, these biological contributions can be neglected. Notwithstanding, there exist models that take these contributions into account [11, 12]. The following derivation is made based on the assumed shape evolution of two spheres as shown in Fig. 3. Initially ($t = 0$) we have two equal sized spheres with radius $a_0$, centered at points $A$ and $B$, connected by one contact point at $O$. During coalescence, both centers gradually move towards $O$. At time $t$, the two intersecting spheres of radius $a(t)$ make a dumb-bell shape. For this configuration, we define the $\theta(t)$ and $x(t)$ as the angle of the intersection and as the radius of the neck, respectively. When coalescence is complete, the center points $A$ and $B$ have merged into point 0 forming a single sphere with radius $a_f$. Combining this shape evolution with mass conservation allows us to write an expression for the radius of the droplet as a function of the angle $\theta$ i.e. the volume of two spheres has to be identical to the volume of two spherical caps

$$\frac{8}{3}\pi a_0^3 = \frac{2\pi}{3}a^3(2 - \cos(\theta))(1 + \cos(\theta))^2. \tag{7}$$

From this we obtain the radius $a(\theta)$

$$a(\theta) = 2^{2/3}a_0(2 - \cos(\theta))^{-1/3}(1 + \cos(\theta))^{-2/3}. \tag{8}$$

The surface area of two spherical caps is calculated by

$$S = 4\pi a(\theta)^2[1+\cos(\theta)] = \frac{8\pi 2^{1/3} a_0^2}{[1+\cos(\theta)]^{1/3}[2-\cos(\theta]^{2/3}}. \tag{9}$$

The work rate of the surface tension $\dot{W}_\Gamma$ is defined by

$$\dot{W}_\Gamma = -\Gamma\frac{dS}{dt} = \Gamma\frac{8\pi a_0^2 2^{1/3}\cos(\theta)\sin(\theta)}{[1+\cos(\theta)]^{4/3}[2-\cos(\theta)]^{5/3}}\dot{\theta}, \tag{10}$$

with $\Gamma$ the surface tension. On the other hand, the viscous dissipation rate for a Newtonian fluid can be described by

$$\dot{W}_\eta = \iiint_V \eta\nabla u : (\nabla u + \nabla u^T)dV, \tag{11}$$

with $\eta$ the viscosity, $\nabla u$ the gradient of velocity, and $V$ the volume of the two coalescing droplets. The flow field is assumed to be extensional and can be described by

$$\nabla u = \begin{bmatrix} \dot{\epsilon}/2 & 0 & 0 \\ 0 & -\dot{\epsilon} & 0 \\ 0 & 0 & \dot{\epsilon}/2 \end{bmatrix} \tag{12}$$

where $\dot{\epsilon}$ is the strain rate. This was a correction of Eshelby on the derivation of Frenkel [13]. From this we find

$$\dot{W}_\eta = \iiint_V 3\eta\dot{\epsilon}^2 dV. \tag{13}$$

Now, according to the original derivation of Frenkel [14], the strain rate $\dot{\epsilon}$ is assumed to be constant throughout the complete domain and can be approximated by

$$\dot{\epsilon} = \frac{\partial u_y(A)}{\partial y} \approx \frac{u_y(A) - u_y(O)}{a}. \tag{14}$$

In this expression, $u_y(O)$ is the velocity of the fluid at the plane of contact of the two particles and is equal to zero. $u_y(A)$ is the velocity at which point A moves towards, i.e., $d/dt[a(\theta)\cos(\theta)]$. Hence, the strain rate is

$$\dot{\epsilon} = \frac{1}{a}\frac{d}{dt}[a(\theta)\cos(\theta)] = \frac{1}{a}[\dot{a}(\theta)\cos(\theta) - a(\theta)\sin(\theta)\dot{\theta}]. \tag{15}$$

Working out these expressions one obtains

$$\frac{dW_\eta}{dt} = 32\pi a_0^3\eta\frac{1-\cos(\theta)}{[1+\cos(\theta)][2-\cos(\theta)]^2}\dot{\theta}^2. \tag{16}$$

Balancing the the work rate of surface tension Eq. (10) with the rate of viscous dissipation Eq. (16) will result in the model by Pokluda [6]. To take into account arrested coalescence, it has been suggested to include an elastic energy term which is caused by internal structure [8, 7]. Following these, the elastic energy can be expressed as

$$W_\varepsilon = \frac{3}{2}G'\epsilon^2 V \tag{17}$$

with

$$\varepsilon = 1 - L(\theta)/L_0 \tag{18}$$

for which they used $L$ as the doublet length Fig. 3. The underlying reasoning is that while surface tension drives the fusion a longitudinal spring tension keeps them apart. In our framework this would imply that

$$\varepsilon = 1 - \frac{a(\theta) + a(\theta)\cos(\theta)}{2a_0}. \tag{19}$$

Before fusion, $a(\theta) = a_0$, there is no strain. When the spheres merge to a full sphere $\theta = 90°$, then $a(\theta) = a_f = 2^{1/3}a_0$. Hence, we find

$$\epsilon(\theta = 90°) = 1 - 2^{-2/3}. \tag{20}$$

Calculating the derivative from Eq. (19) yields

$$\dot{\varepsilon} = -\frac{1}{2a_0}\frac{d}{dt}\left[a(\theta)(1 + \cos(\theta)\right] \tag{21}$$

Note that these elastic strain and strain rate are incompatible with the strain rate described earlier in Eq. (15), following the derivation of Pokluda [6]. However, they are equivalent in magnitude for the onset of fusion i.e. $\theta \approx 0$ and $a(\theta) = a_0$. To preserve the original model in the liquid limit (absence of elasticity), we continue with this hybrid approach. To make this fully transparent in our derivations, we choose to write this strain and strain rate as $\varepsilon$ instead of $\epsilon$. Continuing on Eq. (21), one obtains

$$\dot{\epsilon} = \frac{2^{2/3}\sin(\theta)}{2(1 + \cos(\theta))^{2/3}(2 - \cos(\theta))^{4/3}}\dot{\theta} \tag{22}$$

The elastic energy can be written as $3/2G'\epsilon^2 V$. The work rate of elastic energy under volume conservation assumption is thus defined by

$$\dot{W}_\varepsilon = 3G'\varepsilon\dot{\varepsilon}V \tag{23}$$

After expanding and simplifying all terms we find

$$\frac{dW_\varepsilon}{dt} = 2^{5/3}\pi G' a_0^3 \left[\frac{2(2 - \cos(\theta))^{1/3} - 2^{2/3}(1 + \cos(\theta))^{1/3}}{(2 - \cos(\theta))^{5/3}(1 + \cos(\theta))^{2/3}}\right]\sin(\theta)\dot{\theta} \tag{24}$$

For the other energy terms we can find , based on [6], the surface energy rate

$$\frac{dW_\Gamma}{dt} = \Gamma\frac{8\pi a_0^2 2^{1/3}\cos(\theta)\sin(\theta)}{[1 + \cos(\theta)]^{4/3}[2 - \cos(\theta)]^{5/3}}\dot{\theta}, \tag{25}$$

and viscous dissipation rate:

$$\frac{dW_\eta}{dt} = 32\pi a_0^3\eta\frac{1 - \cos(\theta)}{[1 + \cos(\theta)][2 - \cos(\theta)]^2}\dot{\theta}^2. \tag{26}$$

Now based on conservation of energy; the energy released from lowering the surface area is partially converted to heat by viscous dissipation and partially converted to internal elastic energy i.e. $dW_\Gamma/dt = dW_\eta/dt + dW_\epsilon/dt$. This yields the following differential equation

$$\begin{aligned}\dot{\theta} = &\frac{\Gamma}{a_0\eta}\frac{2^{-5/3}\cos(\theta)\sin(\theta)[2-\cos(\theta)]^{1/3}}{[1+\cos(\theta)]^{1/3}[1-\cos(\theta)]} \\ &-\frac{G'2^{-10/3}}{\eta}\frac{\sin(\theta)}{1-\cos(\theta)} \\ &\left[2(2-\cos(\theta))^{2/3}(1+\cos(\theta))^{1/3} - 2^{2/3}(2-\cos(\theta))^{1/3}(1+\cos(\theta))^{2/3}\right].\end{aligned} \tag{27}$$

In the absence of elasticity $G' = 0$, we trivially retrieve the model as derived by Pokluda [6]. Because this equation is complicated by geometrical factors, we will explore two approximations that further explain the predictions made by this model. First, for small angles we can find

$$\dot{\theta} = \frac{1}{2}\frac{\Gamma}{\eta a_0 \theta} - \frac{1}{8}\frac{G'\theta}{\eta} = \frac{1}{2\tau_\Gamma\theta} - \frac{\theta}{2\tau_\epsilon} = \frac{\tau_\varepsilon - \tau_\Gamma\theta^2}{2\tau_\Gamma\tau_\varepsilon\theta}. \tag{28}$$

Based on this equation we define $\tau_\Gamma := a_0\eta/\Gamma$ and$\tau_\varepsilon := 4\eta/G'$. The first term in this equation is known as the Frenkel-Eshelby model. Fig. 4A&B shows how the two terms of Eq. (27) are approximated by the simple expression of Eq. (28) Solving this differential equation with the initial conditions, $\theta = \theta_0$ for $t = 0$, yields

$$\begin{aligned}\theta &= \sqrt{\frac{\tau_\varepsilon}{\tau_\Gamma} - \left(\frac{\tau_\varepsilon}{\tau_\Gamma} - \theta_0^2\right)\exp\left(-\frac{t}{\tau_\varepsilon}\right)} \\ &= \sqrt{\frac{4\Gamma}{G'a_0} - \left(\frac{4\Gamma}{G'a_0} - \theta_0^2\right)\exp\left(-\frac{G'}{4\eta}t\right)}.\end{aligned} \tag{29}$$

Although theory dictates that coalescence starts at zero angle, we add a non-zero initial condition to account for the initial fast cell relaxation that occurs before the tissue relaxation in our tissue spheroid simulation. However, when we look at the theory $\theta_0 = 0$, so we can investigate limit behavior. On the very onset of coalescence $t \approx 0$, or in the limit of low elasticity $\lim_{G'\to 0} : \exp\left(-\frac{G'}{4\eta}t\right) \approx 1 - \frac{G'}{4\eta}t$. We retrieve the Frenkel-Eshelby model $\theta = \sqrt{\frac{\Gamma}{a_0\eta}t}$. In other words, the initial coalescence will always be liquid-like. Elasticity has an effect later on, which slows down and arrests the fusion. When elasticity is strong, on the other hand, the small angle approximation can be used to calculate the equilibrium angle $\theta = \frac{4\Gamma}{G'a_0}$. Fig. 5A compares this analytical model Eq. (29) with the numerical evaluation of Eq. (27) for various values of $\tau_\varepsilon/\tau_\Gamma$. This plot suggests that this approximation is adequate up to an equilibrium angle of 25°. When larger angles are encountered, a better approximation is required. Following [15], the first term of Eq. (27) can be written as

$$\frac{\Gamma}{a_0\eta}\frac{2^{-5/3}\cos(\theta)\sin(\theta)[2-\cos(\theta)]^{1/3}}{[1+\cos(\theta)]^{1/3}[1-\cos(\theta)]}=\frac{\Gamma}{a_0\eta}\frac{a_0\cot(\theta)}{a(\theta)}. \tag{30}$$

Now, since $1 \leq a(\theta)/a_0 \leq 2^{1/3} \approx 1.26$, they approximate by setting $a(\theta) = a_0$. Note that this means that for the dynamics of $\theta$, volume conservation is ignored. In the absence of elasticity the obtained differential equation is easily calculated with result $\cos(\theta) = \exp\left(-\frac{\Gamma t}{2\eta a_0}\right)$. Also using a geometrical function as an approximation for the second term yields the following differential equation

$$\dot{\theta}=\frac{\Gamma}{2a_0\eta}\cot(\theta)-\frac{G'}{8\eta}\sin(\theta)=\frac{\cot(\theta)}{2\tau_\Gamma}-\frac{\sin(\theta)}{2\tau_\varepsilon}. \tag{31}$$

Based on this equation we define $\tau_\Gamma := a_0\eta/\Gamma$ and $\tau_\varepsilon := 4\eta/G'$. Fig. 4A&B show how the two terms of Eq. (27) are approximated by the simple expression of Eq. (31). We first evaluate the equilibrium solution i.e. $\dot{\theta} = 0$.

$$\cos(\theta_{eq})=-\frac{\tau_\varepsilon}{2\tau_\Gamma}\pm\sqrt{\frac{\tau_\varepsilon^2}{4\tau_\Gamma^2}+1}=-\frac{2\Gamma}{Ga_0}\pm\sqrt{\frac{4\Gamma^2}{G^2a_0^2}+1} \tag{32}$$

Because $0 < \theta < \pi/2$, $\cos(\theta) > 0$, and also $\tau_\varepsilon \geq 0$, $\tau_\Gamma > 0$ so only the positive root is physically possible. According to this approximation, fusion always initiates as $\theta_{eq} = 0$ only occurs at infinite values for G'. On the other hand, complete fusion $\theta_{eq} = 90°$ is only possible in the absence of any elasticity. Fig. 4C shows that although this is still a small angle approximation, it is able to predict the equilibrium angle reasonably well in a wide range. The analytical solution to Eq. (31) is given by

$$\cos(\theta)=-\frac{\tau_\varepsilon}{2\tau_\Gamma}+\sqrt{\frac{\tau_\varepsilon^2}{4\tau_\Gamma^2}+1}\tanh\left(\frac{\sqrt{4\tau_\Gamma^2+\tau_\varepsilon^2}}{4\tau_\Gamma\tau_\varepsilon}(t+C)\right), \tag{33}$$

where $C$ can be obtained from the initial conditions $\theta(t = 0) = \theta_0$

$$C=\frac{4\tau_\Gamma\tau_\varepsilon}{\sqrt{4\tau_\Gamma^2+\tau_\varepsilon^2}}\operatorname{arctanh}\left(\frac{2\tau_\Gamma\cos(\theta_0)+\tau_\varepsilon}{\sqrt{4\tau_\Gamma^2+\tau_\varepsilon^2}}\right) \tag{34}$$

Because $\theta_0 < \theta_{eq}$, $C$ is a complex number. Fig. 5B&C illustrate that this analytical expression nicely approximate the numerical result of Eq. (27).

#### 5 General considerations on quantification of fusion dynamics

In many papers on spheroid fusion, the dynamics of spheroid fusion are analyzed by studying $(x/a_0)^2$ with $x$ the neck radius and $a_0$ the the radius of the spheroid before fusion, shown in Fig. 3, according to the following exponential

$$\left(\frac{x}{a_0}\right)^2 = 2^{2/3}\left(1 - \exp\left(-\frac{t}{\tau}\right)\right), \tag{35}$$

with $\tau = 2^{2/3} a_0\eta/\Gamma \approx 1.58 a_0\eta/\Gamma$ [11, 16], $\tau = 1.35 a_0\eta/\Gamma$ [17], or $\tau = 1.9 a_0\eta/\Gamma$ [9]. Thus, there is a clear difference in proportionality constant between $\tau$ and $a_0\eta/\Gamma$. The method of determining this constant is also, to our knowledge, never mentioned. Of course the exact value of $\tau$ is in practice less important compared to trends and differences between experiments. Unfortunately this might give a skewed look when comparing results with different methodologies. To clear up this confusion we first simulated 50 hours fusion experiments at various values of $\tau = a_0\eta/\Gamma$ using the liquid model of Pokluda [6] and fitted Eq. (35). Then we performed linear regression on the obtained values of $\tau_{\text{fit}}$ to estimate the proportionality constant. Fig. 6A illustrates that the exponential function matches the trend in the data fairly well. However, there is a systematic bias at short and longer times, which becomes more apparent for larger values of $\tau$. Based on the linear regression we identified a proportionality constant of 1.906, which is the same as [9]. However, there is a clear correlation in the deviations around the fitted line Fig. 6C.

In many other papers, spheroid fusion dynamics are analyzed by $\sin^2(\theta) = (x/a)^2$ with $x$ the neck radius and $a$ the the radius of the spheroid at each point during fusion, shown in 3, according to following exponential

$$\sin^2(\theta) = \left(\frac{x}{a}\right)^2 = 1 - \exp\left(-\frac{t}{\tau}\right). \tag{36}$$

This expression can be directly derived from the model of Pokluda [6], following [15],

$$\dot{\theta} = \frac{\Gamma}{a_0\eta}\frac{2^{-5/3}\cos(\theta)\sin(\theta)[2-\cos(\theta)]^{1/3}}{[1+\cos(\theta)]^{1/3}[1-\cos(\theta)]} = \frac{\Gamma}{a_0\eta}\frac{a_0\cot(\theta)}{a(\theta)}. \tag{37}$$

Now, since $1 \leq a(\theta)/a_0 \leq 2^{1/3} \approx 1.26$, they approximate by setting $a(\theta) = a_0$. Note that this implies that for the dynamics of $\theta$, volume conservation is ignored. In the absence of elasticity the obtained differential equation is easily calculated with result $\cos(\theta) = \exp(-\frac{\Gamma t}{2\eta a_0})$. When repeating the same procedure as before, the numerical result and the fit are almost identical Fig. 6B . If we then perform linear regression on the $\tau_{\text{fit}}$ vs $\tau$ the proportionality constant is practically one, Fig. 6D. Furthermore, the prediction interval around the curve is narrow. Based on this analysis, we recommend to analyze complete spheroid fusion using $\sin^2(\theta)$ as a metric.

### 6 Model inconsistencies

In the model of coalescence of two spherical droplet it the liquid is assumed to be incompressible. Then, conservation of mass implies conservation of volume. Subsequently, the radius of the spheroid increases monotonically according to

$$a(\theta) = 2^{2/3} a_0 (2 - \cos(\theta))^{-1/3} (1 + \cos(\theta))^{-2/3}. \tag{38}$$

For complete fusion we have $a(\theta = 90) = a_f = 2^{1/3} a_0 \approx 1.26 a_0$ This monotonous increase is not always observed in our simulations, wherein we simulate aggregation and compaction prior to the fusion. For some parameter combination, aggregation is slow due to its glassy nature and the equilibrium radius is still not reached after 24 hours. Therefore, the further compaction influences the fusion process, which can be seen in $a_0/a_f$ ratio, with $a_0$ the spheroid radius before fusion, and $a_f$ the spheroid radius at the end of the simulation Fig. 7A. In this region, the increase in $\sin^2(\theta) = (x/a)^2$ is both the consequence of the increase in $x$ as the decrease in $a$. To ensure that arrested fusion is not the result of continuing compaction, we sample some parameter settings from the region or arrested fusion and perform the fusion simulations with a different initialization. Firstly, the cell are randomly positioned in a sphere with radius equal to the micro-well radius. Secondly, over a window of 5 hours the sphere is shrunk to a radius of $r_{\text{sphere}} = \sqrt{N \langle r \rangle^3}$, resulting in an apparent density of 1. Then, the compressing sphere is removed from the simulation and the tissue spheroid is allowed to relax for 5 hours to remove the extensive stress by expanding. As in the original setup, the two spheroids are then transferred from their separate micro-wells to a larger micro-well in which fusion naturally progresses. Based on these simulation we conclude that we could successfully circumvent further compaction during fusion; however the fusion remained arrested Fig. 7B&C.

[10] Ioan Kosztin, Gordana Vunjak-Novakovic, and Gabor Forgacs. Colloquium: Modeling the dynamics of multicellular systems: Application to tissue engineering. *Reviews of Modern Physics*, 84, 12 2012.

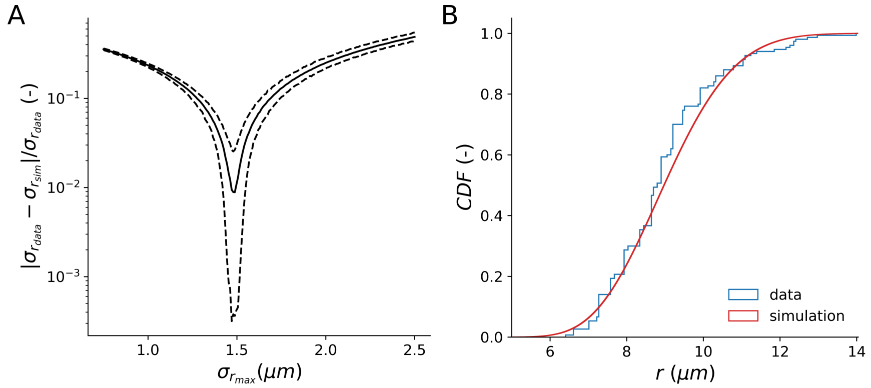


Figure 1: **(A)** Relative error of the simulated standard deviation on the cell radius $\sigma_{\mathrm{r_{max}}}$ with respect to the experimental standard deviation of the cell radius $\sigma_{\mathrm{r_{sim}}}$ expressed a function of $\sigma_{r_{\max}}$. Varying $\sigma_{\mathrm{r_{sim}}} \in [0.75\mu m, 2.5\mu m]$ reveals a distinct minimum in the relative error. After a second finer minimum search we estimated $\sigma_{\mathrm{r_{max}}} = 1.481\mu m$. **(B)** Comparison of simulated probability distribution of cell radii from a simulation of cell growth using the Qu growth model [3] with the distribution of cell radii as experimentally observed in [1].

.

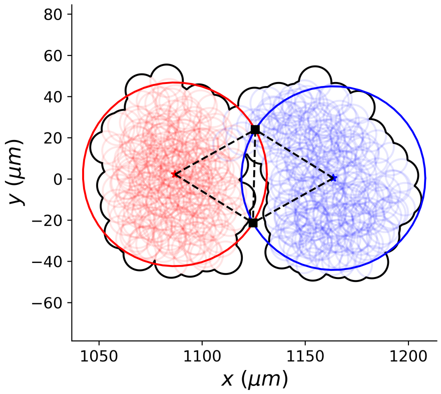


Figure 2: Visualization of shape extraction procedure. Cells (red and blue) are projected in the $X/Y$-plane and the overall contour is computed. Next, two circles of equal radius are fitted to minimize the distance towards the spheroid contours. These circles are used to calculate the effective contact angle $\theta$, see Fig. 3

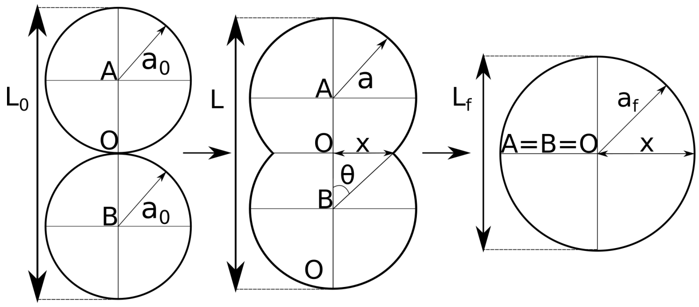


Figure 3: Shape evolution during spheroid fusion adapted form [6]

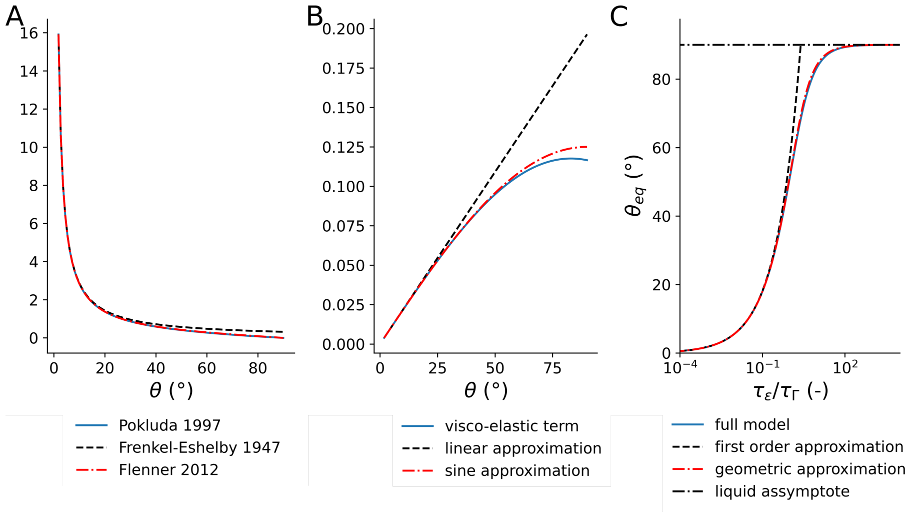


Figure 4: **(A)** Approximation on the visco-capilary term of Eq. (27), which is the fusion model according to Pokluda [6]. While the Frenkel-Eshelby model [14, 13] starts to deviate at larger values of $\theta$, the approximation of Flenner [15] is nearly identical to Pokluda's. **(B)** Approximation on the visco-elastic term of equation Eq. (27). The linear approximation Eq. (28) starts to deviate at angles larger than 25°. The sine approximation follows the trend up to angles of 60°, but the error remains bounded. **(E)** Comparison of the equilibrium angle $\theta_{eq}$ as numerically calculated from Eq. (27) with respect to the first order approximation Eq. (29) and the geometric approximation Eq. (33). Although the geometric approximation is a low angle approximation, but has a bounded error, the trend in equilibrium angle remains consistent with the numerical result. The first order approximation, on the other hand, deviates.

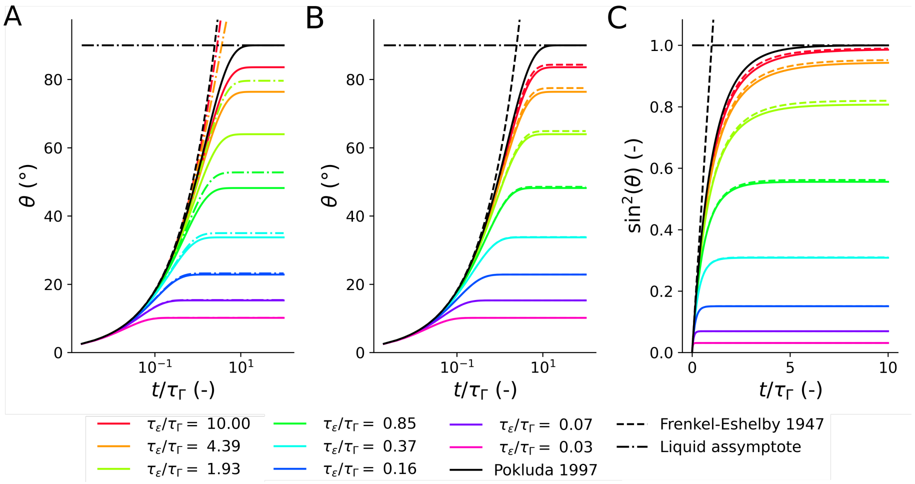


Figure 5: **(A)** Comparison of the numerical result of Eq. (27) (full line) with Eq. (29) (dashed line). As expected this approximation only works for small angles. On the other hand, this graph clearly shows that the initial response is always liquid-like. **(B)** Comparison of the numerical result of Eq. (27) (full line) with Eq. (33)(dashed line). The difference between the two models is slim. **(C)** Comparison of the numerical result of Eq. (27) (full line) with Eq. (33)(dashed line), but shown as $\sin^2(\theta)$, which is often used to analyze the fusion dynamics. This is also how the simulation data will be analyzed. The difference between the two models is slim.

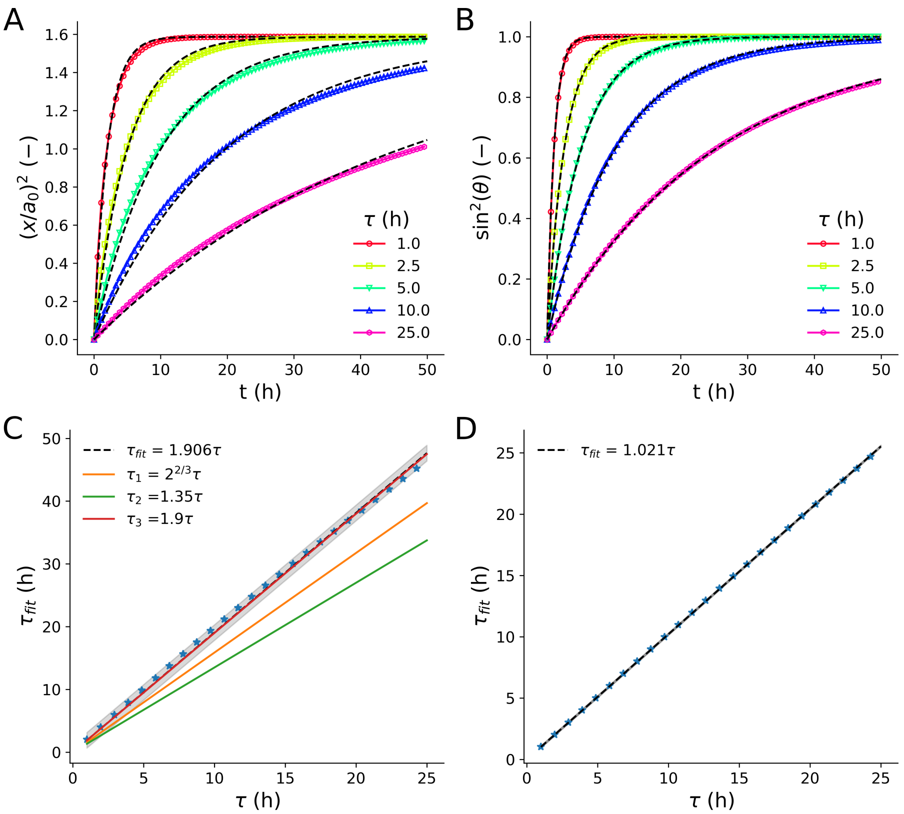


Figure 6: **(A)** and **(B)** show the fusion dynamics simulated using numerical integration of the Pokluda model Eq. (37) expressed as $(x/a_0)^2$ and $\sin^2(\theta) = (x/a)^2$ and fitted by Eq. (35) and by Eq. (36), respectively. In Fig. A, the fit represents the data reasonably well. On the other hand in Fig. B the fit is nearly perfect. **(C)** Relating the simulated time constant with the fitted time constant as done for Fig. A, reveals that they are linearly proportional with factor 1.906. This is in agreement with the one used by Fleming [9]. The error on this regression line is correlated. Also, although this is a deterministic fit, there is some uncertainty, visualized by the prediction interval around the regression line. **(D)** Relating the simulated time constant with the fitted time constant as done for Fig. B, reveals that they are linearly proportional with factor 1.021. In practice this means that there is no need for a correction factor for this approximation. Furthermore, this calibration line fits the data nearly perfect as well.

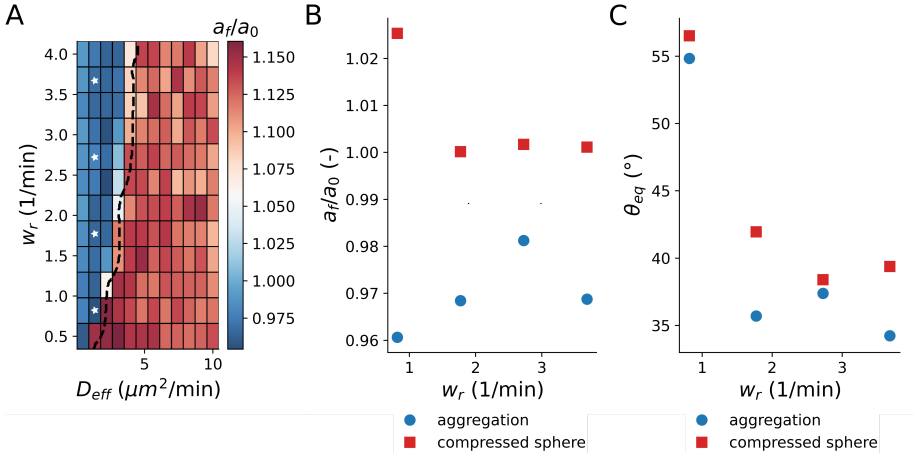


Figure 7: **(A)** Map of the ratio of the average spheroid radius at the end of the simulation $a_f$ to the average spheroid radius before fusion $a_0$ for various values of cell activity $D_{\text{eff}}$ and cell repulsion $w_{\text{r}}$. The black dashed line depicts the boundary between arrested fusion and complete fusion, as obtained from the equilibrium angel $\theta_{eq}$. The region of arrested fusion, overlaps with the zone in which compaction continues during fusion. To verify if this has influence on our classification, four parameter settings from the arrested region were selected, depicted by the stars in the map. **(B)** and **(C)** compare the ratio $a_f/a_0$ and the predicted equilibrium angle $\theta_{eq}$, respectively, when initialization is by aggregation versus compressing sphere procedure for the selected values of $D_{\text{eff}}$ and $w_{\text{r}}$. These graph clearly shows that using the compressing sphere procedure there is no compaction during fusion i.e. $a_f/a_0 > 1$. However, the fusion remained arrested.
